# Cave dogs around major urban areas threaten rabies elimination program

**DOI:** 10.1101/2025.09.28.679082

**Authors:** Ricardo Castillo-Neyra, Elvis W. Díaz, Brinkley Raynor Bellotti, Katherine Morucci, Micaela De la Puente-León, Lizzie Ortiz-Cam, Michael Z. Levy

## Abstract

**Background:** In the city of Arequipa, Peru, the government has implemented control measures against dog rabies virus since the detection of its reintroduction in 2015. The city was previously considered free of animal reservoirs other than owned and stray dogs within its urban boundaries. However, multiple reports from peri-urban residents have suggested the presence of feral dogs living in caves on the city’s outskirts. We aim to document the presence and dietary patterns of feral dogs adjacent to the city margins.

**Methods:** We conducted monthly field visits to four peri-urban localities in eastern Arequipa, an area where the presence of feral dogs had been previously reported. Dog caves were identified by tracking footprints and other field signs left by dogs, and their locations were georeferenced. Each cave was revisited monthly three times to record the presence of live and dead dogs, and puppies. Fecal samples collected around the caves were analyzed to assess dietary patterns.

**Results:** We observed that feral dogs use caves for resting, hiding, and reproduction— some of which appear to be constructed by the dogs themselves. The high number of puppies and dead adult dogs indicates a high population turnover. Dietary analysis revealed that these dogs feed on local fauna, including birds, rodents, cats, sheep, and, notably, other dogs.

**Conclusions:** These unowned, cave-dwelling dogs are not reached by mass rabies vaccination or sterilization programs. Moreover, they exist outside the jurisdiction of health inspectors responsible for rabies surveillance, resulting in a lack of data on rabies infection in this subpopulation. Our findings highlight the need for integrated One Health strategies to address the challenges posed by feral dog populations in rabies elimination efforts.

## INTRODUCTION

The canine rabies elimination program in the city of Arequipa, Peru faces a unique threat that has hitherto been unreported. The geographic positioning of Arequipa amidst the climatically unforgiving Andean high desert was believed to shield the urban fauna (e.g., stray dogs and cats) from contact with local wildlife disease reservoirs (Castillo-Neyra, Brown, et al., 2017). As a result, present rabies control efforts, like those in most Latin American countries (MINSA, 2017; Vigilato et al., 2013), have focused mostly on populations of owned dogs. Control efforts have included a combination of annual, city-wide mass vaccination campaigns, localized vaccination rings around positive cases, passive surveillance of dog populations, as well as methods that have been proven around the world to be ineffective, such as sporadic culling of free-roaming dogs (Castillo Neyra et al., 2016; Morters et al., 2013). Despite these efforts, cases have continued to spread over the past seven years (Raynor et al., 2020). The scope of current control paradigms would be insufficient to address the complexity of the local disease ecology if feral dogs are present in Arequipa.

Feral dog populations have broad negative impacts, from diminishing environmental quality (Joshi et al., 2020; Tebelmann et al., 2025; Young et al., 2011), to displacing native wildlife (Choudhary & Chishty, 2022; Zapata-Ríos & Branch, 2016), to affecting the locals’ livelihoods by targeting livestock (Joshi et al., 2020; Woodroffe et al., 2005).

Most importantly, they pose a significant health risk as potential vectors to a variety of pathogens that affect wildlife, domestic animal species, and human populations (Bergman et al., 2009), such as canine rabies and distemper viruses. Aside from the human impact of rabies and the potential role of feral dogs in the virus transmission, there is also the risk of feral dogs introducing the rabies virus to vulnerable wildlife populations (Bergman et al., 2009; Stuchin et al., 2018; Young et al., 2011), such as the common Andean fox. The establishment of a new native vector and the potential for a novel rabies virus transmission cycle—such as the historical spillover from domestic dogs to red foxes in Europe (Johnston, 2007)—demonstrates how such shifts can complicate efforts to eliminate the disease from a given region.

Recent accounts from peri-urban dwelling inhabitants have detailed numerous sightings and occurrences in which “cave dogs from the hills” have come into the periphery of the city to hunt local domestic animals, including adult pigs and sheep. Herders of medium-sized livestock have reported packs of dogs jumping fences, destroying corrals, and killing farm animals. The presence of feral, cave-dwelling dogs in peri-urban areas on the city’s margins could pose a significant challenge to the rabies control program.

Unowned dogs are neither included in vaccination nor sterilization campaigns, and dogs out of the city boundaries are not monitored by rabies surveillance systems (MINSA, 2017), leaving a critical gap in understanding their role in rabies transmission dynamics. Surveillance and understanding of the behavioral ecology and extent of interactions between populations of owned and unowned dogs in the region are essential to explain the potential contribution of unowned dogs to the local propagation of the canine rabies virus (A.i et al., 1993; Hampson et al., 2009; Wandeler et al., 1988).

In response to these growing concerns, our objectives were to establish the presence of feral dogs in peri-urban areas of Arequipa, characterize their dietary patterns, and explore the threats that these animals present to rabies control programs as well as the One Health challenges that they pose for both local wildlife conservation and the sustainability of small-scale livestock farming.

## MATERIALS AND METHODS

### Study site

The study was conducted in the Alto Selva Alegre (ASA) district, located in the northeastern part of Arequipa, Peru, at an altitude of 2,200 masl. Arequipa city is the second largest city of Peru with a human population of ∼1.1 million. ASA is a medium-sized (∼7 km^2^) and diverse district that includes both densely populated urban areas and less developed peri-urban and natural zones. Its southeastern region borders the foothills of the Misti volcano, a rugged and sparsely inhabited area where sightings and reports of feral or wild dogs have previously occurred. The district shares boundaries with the districts of Cayma, Miraflores, Yanahuara, and Arequipa Cercado.

### Field surveys

Between September and December 2019, we conducted monthly pedestrian surveys within a study area of approximately 1.6 km^2^ located in the northeastern portion of ASA. Our study area comprised three contiguous subdistrict localities—Apsil, El Roble, and San Luis A. The primary goal of these surveys was to identify caves that showed evidence of use by feral dogs and obtain parameters to characterize dogs and caves. A cave was defined as a naturally occurring or canine-modified dwelling in the ground or an earthen wall that was at least 1 meter in-depth, or large enough to permit animal entry and shelter, and also exhibited direct or indirect evidence of dog usage. Caves that were less than a meter in depth, but contained some evidence of dog usage, were georeferenced and recorded as a potential sleeping sight, rather than occupation. In 2022, we revisited the same three localities to collect fecal samples from in and around the identified caves. The goal of this second phase was to perform a dietary analysis to better understand the feeding behavior and ecological role of feral dogs in the area.

### Trails and mapping

During the initial reconnaissance visit, we observed a network of trails formed by the repeated movement of dogs (Figure 1A). This observation informed our decision to explore the use of these trails as a strategy to locate caves. To determine the most effective survey approach, we conducted a short pilot study comparing two methods. In the first, two surveyors followed a predefined grid-based transect, with lines spaced approximately 70 meters apart to form a regular grid. In the second method, two surveyors followed dog-made trails—paths visibly worn by repeated canine use— allowing the natural layout of the trails to guide the search. In one locality, the grid-based transect approach led to the identification of six dog-occupied caves. In contrast, the dog-trail-based approach identified all six of those same caves, along with 17 additional ones, for a total of 23. Based on these results, we selected the dog-trail-based method for the full survey.

**Figure 1.**
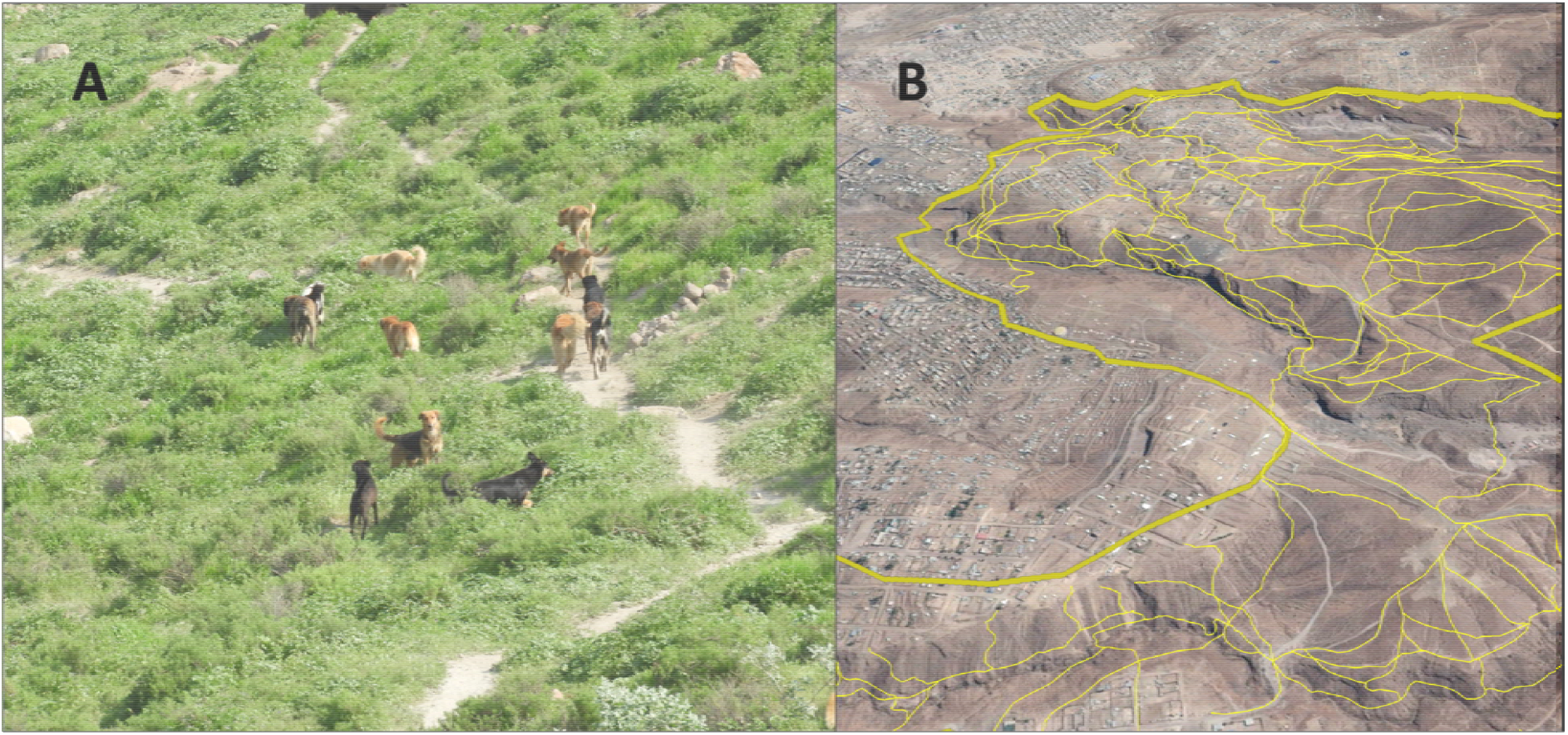
**A:** Pack of dogs in study area leaving a clear path on frequently used tracks. Visible paths were used to determine the transects to search for cave dogs. **B:** The thick yellow line delimits the study area, and thinner yellow lines represent transects followed by our team.

Using Google Earth, we mapped the visible footpaths created by dogs across the study area (Figure 1B) and used these as the basis for our transects. During subsequent visits, paired observers conducted foot surveys along these mapped trails, recording sightings of feral dogs and caves showing evidence of occupation or use.

The region is connected to the rest of the city by a series of semi-natural water channels that serve to collect and deposit rainwater runoff into the city river during the wet season (Castillo-Neyra, Brown, et al., 2017; Raynor et al., 2021). Given the short duration of the wet season, water channels are empty and dry during much of the year and are used by dogs to facilitate movement across city localities and throughout the study region (Castillo-Neyra, Brown, et al., 2017; Raynor et al., 2021). We also visited the water channels and their surroundings in our study area.

### Monthly visitations and data collection

During the first site visitation, we recorded and georeferenced observations of living and dead dogs along with evidence of dog-associated caves. Foot surveys were conducted by a pair of observers along study trails on a monthly basis thereafter.

*Direct evidence* was limited to the sighting of adult or juvenile dogs either in the cave itself or in the direct vicinity. *Indirect evidence* of dog occupation included the presence of dog feces, paw prints, claw marks, and remains of scavenged food. Identified caves included individual dwellings as well as series of interconnected cave structures that occurred within two-meter distance of one another.

All data were recorded using data forms that we developed for the unique purposes of this study on the World Veterinary Service (WVS) mobile phone application (Gibson et al., 2018). These forms were designed to collect and georeference data pertaining to direct sightings of individual live or dead dogs, animal packs, litters, locations, and qualities of associated caves, along with multiple types of indirect evidence as described above. Notably, the WVS app can be used offline, which was valuable considering internet and mobile signals were unavailable throughout the majority of our study area.

### Cave data characterization

We estimated the density of caves stratified by occupancy status and the density of dogs stratified by age and alive status. For each visit, we calculated the mean number of dogs, stratified by age and alive status, as well as the proportion of caves showing signs of occupancy, stratified by the type of indirect evidence observed. We took pictures of observed dogs and reviewed them manually in our lab to prevent double-counting. All data handling and characterization were conducted in R (R Core Team, 2023).

### Dietary analysis

Feces were collected and geolocalized across the study region between March 1st and 4th of 2022. A dietary analysis was performed on 114 samples. Feces were washed in a bowl, and macroscopic content was left out to dry. Samples were then roughly sorted into grossly appreciable categories, including bone, fur, feather, eggshell, claw, plant fiber, unidentifiable organic matter and inorganic matter. Contents in each category were enumerated and skeletal and dental remains were identified by an individual with training in Andean zooarchaeology (K.M.). Prior to analysis, a list of common, morphologically distinct, indigenous, introduced, and domestic fauna in the Alto Selva Alegre District was compiled (Supplement 1) and served as a comparative list for osteological identifications. Based on the quality of digested bone, elements were identified based on representative taxa and skeletal elements. Remains that were highly damaged, fragmented, or otherwise non-diagnostic were labeled as unidentifiable.

## RESULTS

### Cave and dog findings

We identified 158 caves throughout the 1.6 km^2^ study area. Many caves showed claw marks suggesting they were created or expanded by dogs (Figure 2). On average, there were 98.8 caves per square kilometer in the study area, with one locality having 133.3 caves per square kilometer (Table 1). All but 1 cave (99.4%) exhibited either direct or indirect evidence of dog use during each of the consecutive monthly visitations. Furthermore, 7.9% of these caves exhibited direct evidence of their occupation by dogs, meaning that dogs were observed either inside the cave or directly surrounding the cave at the time of the survey.

**Table 1.** Parameters of caves and dogs found in the northeastern outskirts of Arequipa city, Peru.

|  | **Overall** | **APSIL** | **San Luis A** | **El Roble** |
| --- | --- | --- | --- | --- |
| **Study areas** | 1.60 km<sup>2</sup> | 0.73 km<sup>2</sup> | 0.39 km<sup>2</sup> | 0.48 km<sup>2</sup> |
| **Cave density** | 98.75/km<sup>2</sup> | 72.60/km<sup>2</sup> | 133.33/km<sup>2</sup> | 110.42/km<sup>2</sup> |
| **Cave with direct evidence density** | 7.81//km<sup>2</sup> | 10.27//km<sup>2</sup> | 8.97//km<sup>2</sup> | 3.13//km<sup>2</sup> |
| **Cave with indirect evidence density** | 61.25//km<sup>2</sup> | 30.14//km<sup>2</sup> | 103.21//km<sup>2</sup> | 74.48//km<sup>2</sup> |
| **Mean live adult dogs per visit [range]** | 32.0 [23 - 41] | 19.00 [15 - 25] | 3.50 [1 - 5] | 9.50 [1 - 21] |
| **Mean dead dogs per visit [range]** | 13.75 [9 - 18] | 7.75 [4 - 10] | 1.25 [1 - 2] | 4.75 [2 - 8] |
| **Mean litters per visit [range]** | 4.25 [2 - 7] | 3.75 [1 - 6] | 0.25 [0 - 1] | 0.25 [0 - 1] |
| **Mean puppies per visit [range]** | 13.75 [7 - 23] | 11.25 [1 - 19] | 1.00 [0 - 4] | 1.50 [0 - 6] |

**Figure 2.**
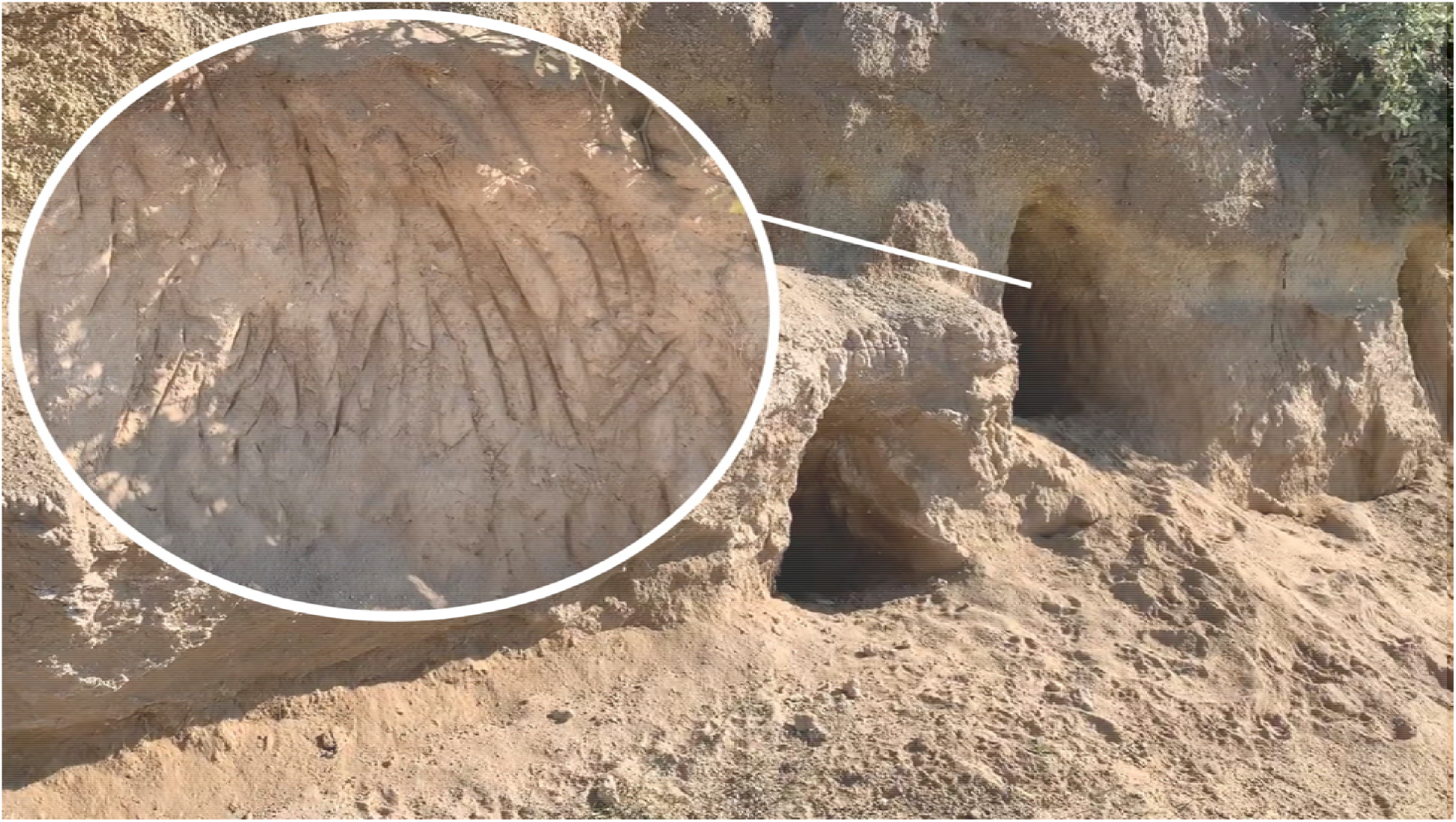
Groups of dog caves in a hill within the study area. Inset shows claw marks on the cave walls.

Over the course of four monthly visits, we observed a total of 128 live adult dogs, 55 dead adult dogs, and 17 litters comprising 55 puppies. Fieldworkers noted that the vast majority of the puppies appeared to be female, although individual-level data on sex were not systematically recorded. On average per visit, we observed 32 live adult dogs, 13.7 puppies, and 13.7 dead adult dogs, with monthly counts varying across visits (Table 1). Notably, some dead dogs were found in tight spatial clusters. Litters were detected in two of the three study localities during each round of data collection. Surprisingly, no dead puppies were observed during any of the four visits.

### Dietary Analysis

We collected 194 fecal samples from various caves. Recovered materials included beaks, bones, claws, feathers, plant fibers, keratin, plastic fragments, eggshells, skin, and teeth (Table 2). Most identifiable elements consisted of skeletal remains. Of the 100 bones recovered, 43 were unidentifiable. The remaining 57 were classified to the following taxa: birds (25/57; 44%), dogs (21/57; 37%), sheep (5/57; 9%), rodents (4/57; 7%), and cats (2/57; 3%). Within the bird and rodent categories, some remains were consistent with domestic species such as chickens and guinea pigs—common backyard livestock in the area. Other specimens were suggestive of additional wild species, but conclusive identification was not possible.

**Table 2.** Dietary analysis from feces collected in and around caves inhabited by feral dogs in Arequipa, Peru, in 2022.

| **Material** | **Count (n=194)** | **Percent** | **95% CI** |
| --- | --- | --- | --- |
| Bone | 100 | 51.55% | 44.28 - 58.76 |
| Claw | 27 | 13.92% | 9.38 - 19.60 |
| Fiber | 26 | 13.40% | 8.95 - 19.02 |
| Feather | 18 | 9.28% | 5.59 - 14.27 |
| Teeth | 8 | 4.12% | 1.80 - 7.96 |
| Eggshell | 7 | 3.61% | 1.46 - 7.29 |
| Skin | 5 | 2.58% | 0.84 - 5.91 |
| Beak | 1 | 0.52% | 0.01 - 2.84 |
| Keratin | 1 | 0.52% | 0.01 - 2.84 |
| Plastic | 1 | 0.52% | 0.01 - 2.84 |

## DISCUSSION

Our field investigations confirm the presence of feral, cave-dwelling dog populations in peri-urban areas bordering the city of Arequipa, Peru—an observation consistent with previous reports in Arequipa, and a problem that has been reported in other countries in the Americas (Bergman et al., 2009; Tebelmann et al., 2025; Zanini et al., 2023; Zapata-Ríos & Branch, 2016). Here, we provide new quantitative, demographic, and dietary data that substantiate and expand upon those findings. We identified feral dog populations inhabiting caves in the southeastern peri-urban outskirts of Arequipa, particularly in Alto Selva Alegre District. Following the survey presented in this study, we conducted exploratory visits to three additional districts surrounding the city and observed similar conditions—hundreds of caves inhabited by feral dogs. The presence of these dogs challenges previous assumptions that urban dog populations are largely isolated from wildlife disease reservoirs and unmanaged canine populations (Castillo-Neyra, Zegarra, et al., 2017; Raynor et al., 2020).

Our dietary analysis of 194 fecal samples revealed evidence of a varied and opportunistic diet. Almost half of all recovered skeletal elements were unidentifiable—a proportion consistent with expected bone degradation from digestion and gnawing (Klare et al., 2011). The avian remains likely include domestic chickens, as suggested by the size and morphology of claw sheaths, skin, and bones. Informal conversations with local residents further support the notion that feral dogs hunt backyard livestock in addition to scavenging from household garbage (Contreras-Abarca et al., 2022; Duarte et al., 2016; Gonzaga et al., 2024), which is frequently discarded into dry riverbeds due to lack of formal waste management. Notably, canine skeletal remains comprised over one-third of all identifiable bones. Several fecal samples contained phalanges and metapodial elements from medium-to large-sized dogs, suggesting possible scavenging or intra-species predation—behaviors that may indicate nutritional stress or limited food availability. Although local accounts describe feral dogs hunting cats and other dogs in packs, further behavioral observation is needed to clarify whether these animals primarily scavenge (Joshi et al., 2020; Meyer et al., 2003) or actively prey on conspecifics (Daniels, 1987)and other species.

Canine rabies remains endemic in the city of Arequipa despite ongoing control and elimination efforts (Bellotti et al., 2025; Xie et al., 2025). Although suboptimal vaccination coverage among owned dogs is the primary factor sustaining transmission (Castillo-Neyra et al., 2019), unrecognized or unsurveyed subpopulations—such as feral, cave-dwelling dogs—may also contribute to the persistence of the virus (Bergman et al., 2009). As the region approaches elimination thresholds, the relative importance of these overlooked groups is likely to increase. Rabies elimination strategies hinge on achieving and maintaining 70% vaccination coverage across the entire at-risk canine population (WHO, 2013). However, most vaccination campaigns are designed to reach owned dogs within established administrative boundaries, potentially excluding adjacent feral populations that remain unvaccinated and unsurveyed (MINSA, 2017).

Our findings underscore the spatial and ecological connectivity between urban, peri-urban, and rural dog populations, highlighting the need for expanded surveillance and control efforts that include feral dog groups. The presence of these unsurveyed populations may undermine progress toward regional and global rabies elimination goals (WHO et al., 2018). While the expansion of rabies in southern Peru has largely been attributed to human migration and the translocation of owned dogs (Salazar et al., 2025), the widespread presence of feral dogs across the Arequipa region suggests that their potential role in disease persistence and spread warrants consideration. Notably, among the observed litters, the vast majority of puppies were female—an unusual deviation from the expected 1:1 sex ratio. Informal conversations with local residents indicated that people, including dog vendors, frequently visit the area to take male puppies, either for sale in local markets—where male dogs are preferred—or for use as guard animals. This ongoing removal and movement of unvaccinated puppies from unmonitored populations into the city presents an overlooked risk that should be considered in the design and implementation of rabies control programs.

In addition to their potential role in rabies epidemiology, the presence of these dogs has other important implications for public health. Settlements within a few hundred meters of these caves report some of the highest rates of dog bites globally, yet display low rates of post-exposure prophylaxis and healthcare-seeking behavior (De la Puente-León et al., 2020). While residents do not distinguish between city dogs and feral cave dogs in reports of bites, the spatial proximity and possible movement between these populations complicate surveillance and control efforts.

Worldwide, feral dogs also pose threats to local economies and biodiversity. Their presence is often first reported through farmers’ and herders’ complaints about livestock attacks (Contreras-Abarca et al., 2022; Duarte et al., 2016; Gonzaga et al., 2024). In ASA and surrounding districts, small-scale animal husbandry remains a key economic activity. Repeated predation on domestic animals undermines household income and food security. Moreover, wildlife authorities have documented feral dog predation on protected species, including South American camelids and Andean condors (Monar-Barragán et al., 2023; Silva Rochefort & Root-Bernstein, 2021). The potential for interspecific competition and disease transmission with native carnivores such as the Andean fox is a further concern. Feral dogs may act as reservoirs for pathogens such as canine distemper virus and sarcoptic mange, both of which can cause substantial mortality in wild canid populations (Acebes et al., 2022; Beineke et al., 2015).

Addressing the ecological and public health impacts of feral dog populations requires cross-sectoral strategies—an approach that is progressing slowly in Arequipa (Castillo-Neyra et al., 2025). Surveillance programs targeting cave-dwelling dogs are essential to understanding their movements, population dynamics, and role in zoonotic disease transmission. Sterilization and vaccination efforts could help reduce their ecological impact and interrupt rabies transmission chains. However, high population turnover remains a significant barrier to the long-term effectiveness of these interventions. Importantly, this challenge is not limited to feral populations: high turnover is also observed among owned dogs in Arequipa, particularly in periurban communities (Diaz et al., 2025). In these areas, vaccinated and sterilized feral dogs may be rapidly replaced by unvaccinated, intact dogs—often due to abandonment from nearby periurban neighborhoods. Establishing the prevalence of infectious diseases such as rabies, distemper, and mange in these feral and periurban owned dog populations will inform both wildlife conservation efforts and the design of effective public health interventions (Morters et al., 2014; Prager et al., 2012; Schurer et al., 2014). Ultimately, controlling rabies and mitigating other health and environmental risks posed by feral dogs in peri-urban Arequipa will require adaptive, integrated One Health strategies that account for the complex ecological interfaces between humans, domestic animals, and wildlife.

## Acknowledgments

This project was supported by NIH-NIAID (K01AI139284 to RCN) and the Fogarty International Center (D43TW012741 to RCN, EWD, and LOC). The funders had no role in study design, data collection and analysis, decision to publish, or preparation of the manuscript.

## REFERENCES

Acebes, P., Vargas, S., & Castillo, H. (2022). Sarcoptic mange outbreaks in vicuñas (Cetartiodactyla: Camelidae): A scoping review and future prospects. Transboundary and Emerging Diseases, 69(5), e1201–e1212. 10.1111/tbed.14479

Ai, W., H.c, M., A, K., & A, B. (1993). The ecology of dogs and canine rabies: A selective review. 12(1), 51. 10.20506/rst.12.1.663

Beineke, A., Baumgärtner, W., & Wohlsein, P. (2015). Cross-species transmission of canine distemper virus—An update. One Health, 1, 49–59. 10.1016/j.onehlt.2015.09.002

Bellotti, B. R., Díaz, E. W., Puente-León, M.D.la, Rieders, M. T., Recuenco, S. E., Levy, M. Z., & Castillo-Neyra, R. (2025). Challenging a paradigm: Staggered versus single-pulse mass dog vaccination strategy for rabies elimination. PLOS Computational Biology, 21(2), e1012780. 10.1371/journal.pcbi.1012780

Bergman, D., Breck, S. W., & Bender, S. (2009). Dogs Gone Wild: Feral Dog Damage in the United States. 8.

Castillo Neyra, R., Levy, M. Z., & Náquira, C. (2016). Effect of free-roaming dogs culling on the control of canine rabies | Revista Peruana de Medicina Experimental y Salud Pública. https://rpmesp.ins.gob.pe/index.php/rpmesp/article/view/2564

Castillo-Neyra, R., Brown, J., Borrini, K., Arevalo, C., Levy, M. Z., Buttenheim, A., Hunter, G. C., Becerra, V., Behrman, J., & Paz-Soldan, V. A. (2017). Barriers to dog rabies vaccination during an urban rabies outbreak: Qualitative findings from Arequipa, Peru. 10.1371%2Fjournal.pntd.0005460

Castillo-Neyra, R., Ortiz-Cam, L., Cañari-Casaño, J. L., Diaz, E. W., Tamayo, L. D., Porras, G., Recuenco, S. E., & Paz-Soldan, V. A. (2025). An Implementation Science Framework to Understand Low Coverage in Mass Dog Rabies Vaccination. medRxiv: The Preprint Server for Health Sciences, 2025.02.05.25321168. 10.1101/2025.02.05.25321168

Castillo-Neyra, R., Toledo, A. M., Arevalo-Nieto, C., MacDonald, H., De la Puente-León, M., Naquira-Velarde, C., Paz-Soldan, V. A., Buttenheim, A. M., & Levy, M. Z. (2019). Socio-spatial heterogeneity in participation in mass dog rabies vaccination campaigns, Arequipa, Peru. PLoS Neglected Tropical Diseases, 13(8), e0007600. 10.1371/journal.pntd.0007600

Castillo-Neyra, R., Zegarra, E., Monroy, Y., Bernedo, R. F., Cornejo-Rosello, I., Paz-Soldan, V. A., & Levy, M. Z. (2017). Spatial Association of Canine Rabies Outbreak and Ecological Urban Corridors, Arequipa, Peru. Tropical Medicine and Infectious Disease, 2(3), 38. 10.3390/tropicalmed2030038

Choudhary, N., & Chishty, N. (2022). Impact of Feral Dogs on Wildlife Community.

Contreras-Abarca, R., Crespin, S. J., Moreira-Arce, D., & Simonetti, J. A. (2022). Redefining feral dogs in biodiversity conservation. Biological Conservation, 265, 109434. 10.1016/j.biocon.2021.109434

Daniels, T. J. (1987). Conspecific Scavenging by a Young Domestic Dog. Journal of Mammalogy, 68(2), 416–418. 10.2307/1381488

De la Puente-León, M., Levy, M. Z., Toledo, A. M., Recuenco, S., Shinnick, J., & Castillo-Neyra, R. (2020). Spatial Inequality Hides the Burden of Dog Bites and the Risk of Dog-Mediated Human Rabies. The American Journal of Tropical Medicine and Hygiene, 103(3), 1247–1257. 10.4269/ajtmh.20-0180

Diaz, E. W., Sila, S., Raynor, B. B., Puente-León, M.D.la, Levy, M. Z., & Castillo-Neyra, R. (2025). Highly Dynamic Population of Owned Dogs and Implications for Zoonosis Control (p. 2025.04.23.650047).bioRxiv. 10.1101/2025.04.23.650047

Duarte, J., García, F. J., & Fa, J. E. (2016). Depredatory impact of free-roaming domestic dogs on Mediterranean deer in southern Spain: Implications for human-wolf conflict. Folia Zoologica, 65(2), 135–141. 10.25225/fozo.v65.i2.a8.2016

Gibson, A. D., Mazeri, S., Lohr, F., Mayer, D., Bailey, J. L. B., Wallace, R. M., Handel, I. G., Shervell, K., Bronsvoort, B. M. deC, Mellanby, R. J., & Gamble, L. (2018). One million dog vaccinations recorded on mHealth innovation used to direct teams in numerous rabies control campaigns. PLOS ONE, 13(7), e0200942. 10.1371/journal.pone.0200942

Gonzaga, M. da C., Borges, J. R. J., Alves, T. S., de Sousa, D. E. R., de Castro, M. B., & Câmara, A. C. L. (2024). Domestic dog attacks on livestock referred to a Veterinary Teaching Hospital. Frontiers in Veterinary Science, 11, 1342258. 10.3389/fvets.2024.1342258

Hampson, K., Dushoff, J., Cleaveland, S., Haydon, D. T., Kaare, M., Packer, C., & Dobson, A. (2009). Transmission Dynamics and Prospects for the Elimination of Canine Rabies. PLoS Biology, 7(3), e1000053. 10.1371/journal.pbio.1000053

Johnston, D. H. (2007). Historical Perspective of Rabies in Europe and the Mediterranean Basin. The Canadian Veterinary Journal, 48(5), 535. https://www.ncbi.nlm.nih.gov/pmc/articles/PMC1852597/

Joshi, B. D., Sharma, L. K., Kaur, A., & Chandra, K. (2020). Assessment of population and impacts of feral dogs on wildlife livestock and humans to design a feral dog management strategy in the Lahaul-Pangi landscape of Himachal Pradesh. https://hpforest.gov.in/storage/files/4/aboutus/wildlifefile15-07-2023-1689415376.pdf

Klare, U., Kamler, J. F., & MacDonald, D. (2011). A comparison and critique of different scatUanalysis methods for determining carnivore diet. https://onlinelibrary.wiley.com/doi/10.1111/j.1365-2907.2011.00183.x

Meyer, W., Schnapper, A., & Eilers, G. (2003). Garbage-dependent nutrition of wild canids and stray dogs: Part 2: Stray dogs and dangers. https://www.researchgate.net/publication/288628927_Garbage-dependent_nutrition_of_wild_canids_and_stray_dogs_Part_2_Stray_dogs_and_dangers

MINSA. (2015). Boletín Epidemiológico (Lima). https://www.dge.gob.pe/portal/docs/vigilancia/boletines/2015/15.pdf

MINSA. (2017). Norma técnica de salud, para la vigilancia, prevención y control de la rabia humana en el Perú. http://bvs.minsa.gob.pe/local/MINSA/4193.pdf

Monar-Barragán, P., Araujo, E., Restrepo-Cardona, J., Kohn, S., Paredes-Bracho, A., & Vargas, F. (2023). Impacts of Free-Ranging Dogs on a Community of Vertebrate Scavengers in a High Andean Ecosystem. Tropical Conservation Science, 16, 1– 10. 10.1177/19400829231218409

Morters, M. K., McKinley, T. J., Restif, O., Conlan, A. J. K., Cleaveland, S., Hampson, K., Whay, H. R., Damriyasa, I. M., & Wood, J. L. N. (2014). The demography of free□roaming dog populations and applications to disease and population control. https://besjournals.onlinelibrary.wiley.com/doi/10.1111/1365-2664.12279

Morters, M. K., Restif, O., Hampson, K., Cleaveland, S., Wood, J. L. N., & Conlan, A. J.K. (2013). Evidence-based control of canine rabies: A critical review of population density reduction. Journal of Animal Ecology, 82(1), 6–14. 10.1111/j.1365-2656.2012.02033.x

Prager, K. C., Mazet, J. A. K., Munson, L., Cleaveland, S., Donnelly, C. A., Dubovi, E. J., Szykman Gunther, M., Lines, R., Mills, G., Davies-Mostert, H. T., Weldon McNutt, J., Rasmussen, G., Terio, K., & Woodroffe, R. (2012). The effect of protected areas on pathogen exposure in endangered African wild dog (Lycaon pictus) populations. Biological Conservation, 150(1), 15–22. 10.1016/j.biocon.2012.03.005

R Core Team. (2023). R: A Language and Environment for Statistical Computing. https://www.r-project.org/

Raynor, B., De la Puente-León, M., Johnson, A., Díaz, E. W., Levy, M. Z., Recuenco, S. E., & Castillo-Neyra, R. (2020). Movement patterns of free-roaming dogs on heterogeneous urban landscapes: Implications for rabies control. Preventive Veterinary Medicine, 178, 104978. 10.1016/j.prevetmed.2020.104978

Raynor, B., Diaz, E. W., Shinnick, J., Zegarra, E., Monroy, Y., Mena, C., De la Puente-León, M., Levy, M. Z., & Castillo-Neyra, R. (2021). The impact of the COVID-19 pandemic on rabies reemergence in Latin America: The case of Arequipa, Peru. 10.1371/journal.pntd.0009414

Salazar, R., Brunker, K., Díaz, E. W., Zegarra, E., Monroy, Y., Baldarrago, G. N., Borrini-Mayorí, K., Puente-León, M.D.la, Palmalux, N., Nichols, J., Kasaragod, S., Levy, M. Z., Hampson, K., & Castillo-Neyra, R. (2025). Genomic characterization of a dog-mediated rabies outbreak in El Pedregal, Arequipa, Peru. PLOS Neglected Tropical Diseases, 19(3), e0012396. 10.1371/journal.pntd.0012396

Schurer, J. M., Phipps, K., Okemow, C., Beatch, H., & Jenkins, E. (2014). Stabilizing Dog Populations and Improving Animal and Public Health Through a Participatory Approach in Indigenous Communities. Zoonoses and Public Health, 62. 10.1111/zph.12173

Silva Rochefort, B., & Root-Bernstein, M. (2021). History of canids in Chile and impacts on prey adaptations. https://onlinelibrary.wiley.com/doi/full/10.1002/ece3.7642

Stuchin, M., Machalaba, C. M., Olival, K. J., Artois, M., Bengis, R. G., Caceres, P., Diaz, F., Erlacher-Vindel, E., Forcella, S., Leighton, F. A., Murata, K., Popovic, M., Tizzani, P., Torres, G., & Karesh, W. B. (2018). Rabies as a threat to wildlife. Revue Scientifique Et Technique (International Office of Epizootics), 37(2), 341– 357. 10.20506/rst.37.2.2858

Tebelmann, H., Kaefer, S., Pages, A., Ruiz, A., Volkart, N., & Ganslosser, U. (2025). Dogs gone wild: Habitat use and ecological impacts of feral dogs in sub-Antarctic Chile. 10.1101/2025.05.08.652634

Vigilato, M. A. N., Clavijo, A., Knobl, T., Silva, H. M. T., Cosivi, O., Schneider, M. C., Leanes, L. F., Belotto, A. J., & Espinal, M. A. (2013). Progress towards eliminating canine rabies: Policies and perspectives from Latin America and the Caribbean. Philosophical Transactions of the Royal Society of London. Series B, Biological Sciences, 368(1623), Article 1623. 10.1098/rstb.2012.0143

Wandeler, A. I., Budde, A., Capt, S., Kappeler, A., & Matter, H. (1988). Dog Ecology and Dog Rabies Control. Reviews of Infectious Diseases, 10(Supplement_4), S684–S688. 10.1093/clinids/10.Supplement_4.S684

WHO. (2013). WHO Expert Consultation on Rabies. Second Report. https://www.who.int/publications/i/item/WHO-TRS-982

WHO, F.O. & OIE. (2018). Zero by 30: The global strategic plan to end human deaths from dog-mediated rabies by 2030. https://www.woah.org/fileadmin/Home/eng/Media_Center/docs/Zero_by_30_FINAL_online_version.pdf

Woodroffe, R., Lindsey, P., Romañach, S., Stein, A., & ole Ranah, S. M. K. (2005). Livestock predation by endangered African wild dogs (Lycaon pictus) in northern Kenya. Biological Conservation, 124(2), 225–234. 10.1016/j.biocon.2005.01.028

Xie, S., Shinnick, J., Diaz, E. W., Zegarra, E., Monroy, Y., Recuenco, S. E., & Castillo-Neyra, R. (2025). Neighborhood Socioeconomic Status and Dog-Mediated Rabies: Disparities in Incidence and Surveillance Effort in a Latin American City (p. 2025.04.02.25325110). medRxiv. 10.1101/2025.04.02.25325110

Young, J. K., Olson, K. A., Reading, R. P., Amgalanbaatar, S., & Berger, J. (2011). Is Wildlife Going to the Dogs? Impacts of Feral and Free-roaming Dogs on Wildlife Populations. BioScience, 61(2), 125–132. 10.1525/bio.2011.61.2.7

Zanini, F., Di Salvo, V., Pierangeli, N., Lazzarini, L., & Curto, E. (2023). Presence of Echinococcus granulosus sensu lato in the endoparasitic fauna of feral dogs in Tierra del Fuego, Argentina. Veterinary Parasitology: Regional Studies and Reports, 44, 100916. 10.1016/j.vprsr.2023.100916

Zapata-Ríos, G., & Branch, L. C. (2016). Altered activity patterns and reduced abundance of native mammals in sites with feral dogs in the high Andes. Biological Conservation, 193, 9–16. 10.1016/j.biocon.2015.10.016

